# MiR-138-5p upregulation during neuronal maturation parallels with an increase in neuronal survival

**DOI:** 10.1101/2022.11.15.516547

**Authors:** M. Asunción Barreda-Manso, Altea Soto, Teresa Muñoz-Galdeano, David Reigada, Manuel Nieto-Díaz, Rodrigo M. Maza

## Abstract

Neuronal maturation is a process that plays a key role in the development and regeneration of the central nervous system. Although embryonic brain development and neurodegeneration have received considerable attention, the events that govern postnatal neuronal maturation are less understood. Among the mechanisms influencing such neuronal maturation process, apoptosis plays a key role. Several regulators have been described to modulate apoptosis, including post-transcriptional regulation by microRNAs. This study aimed to assess whether the strikingly induced miR-138-5p during neuronal maturation contributes to avoiding the neuronal death induced by apoptosis. Our results point out that the observed opposite expression of miR-138-5p and its target Caspase3 might modulate apoptosis favouring neuronal survival at distinct maturation stages. The unchanged expression of miR-138-5p in mature neurons contrasts with the significant downregulation in immature neurons upon apoptotic stimulation. Similarly, immunoblot and individual cellular assays confirmed that during maturation, not only the expression but processing of CASP-3 and caspase activity is reduced after apoptotic stimulation which resulted in a reduction of neuronal death. For all this data, we suggest that the upregulation of miR-138-5p during neuronal maturation is crucial in neuronal survival in pathological or traumatic conditions. Further studies would be needed to determine a more detailed role of miR-138-5p in apoptosis during neuronal maturation and the synergistic action of several microRNAs acting cooperatively on Caspase3 or other apoptotic targets.

**Graphical abstract:** 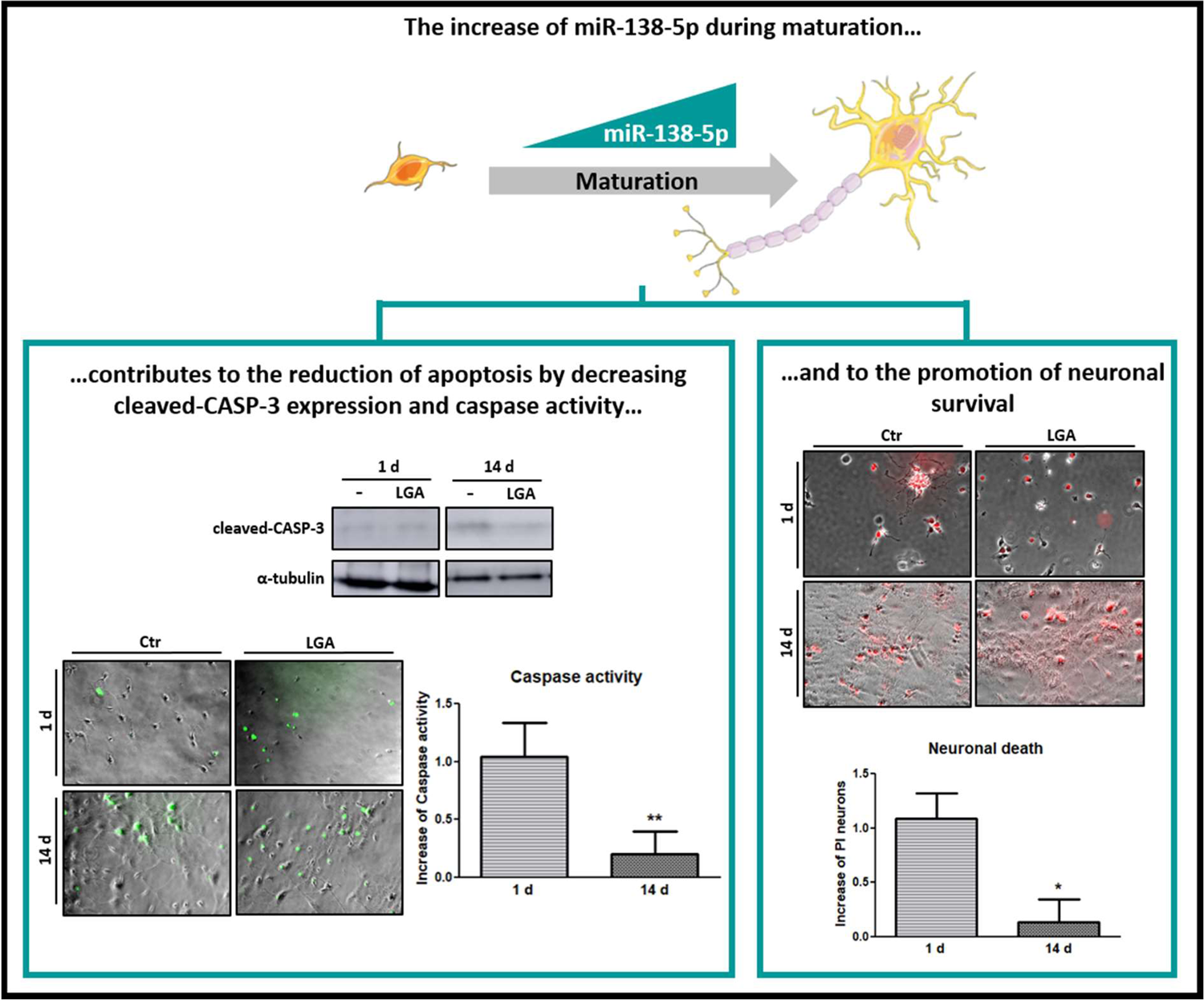

**Highligths:**

- Neuronal maturation promotes cell survival.
- Neuronal maturation reduces the processing of CASP-3 and caspase activity after apoptotic stimulation.
- The expression changes of miR-138-5p and Caspase 3 are opposite during hippocampal neuron maturation.
- The increase of miR-138-5p expression throughout maturation is not influenced by apoptotic stimulation.
- MiR-138-5p is suggested as an essential factor concerning neuronal maturation, regulating Caspase 3 and favouring survival.

## 1. Background

Neurogenesis is the process of formation of new neurons and involves the proliferation, migration, differentiation, and integration of neurons into the existing neuronal circuit. Within these 4 neurogenesis phases, neuronal maturation plays a primordial role, which is not only relevant during the development of the individual, but also in the creation of new neurons in the adult individual for essential processes such as memory [Aimone *et al*., 2006], or even during neurodegenerative diseases or injury repair [Ming, 2011]. Because of this, neuronal maturation has a strong impact on the understanding of the central nervous system (CNS) functioning, in both physiological and pathological conditions. Therefore, it is essential to study the mechanisms of action underlying this maturation, in order to promote it and thus achieve the necessary regeneration in many pathological and traumatic processes [Bradke *et al*., 2020].

Neuronal maturation is a complex process in which, over days and even weeks, neurons change morphologically, including axonal growth, dendrite formation, and connection of the dendritic tree with neighbouring neurons [Piatti *et al*., 2006]. Various factors cause these changes, such as environment [Piatti *et al*., 2011; Tooley *et al*., 2021], growth factors (e.g. NGF) [Crutcher, 1986; Landreth, 1999; Aloe *et al*., 2015], and transcriptome [He and Yu, 2018; Patel *et al*., 2021]. Modulation of transcriptome by non-coding RNA includes microRNAs, small RNA molecules of about 19-25 nucleotides capable of controlling the expression of hundreds of genes and regulating the state and fate of cells [Kosik, 2010]. MicroRNAs have a pivotal role throughout neuronal development and maturation, from the formation of different neuronal types to the decision of neuronal plasticity and connections with other neurons [Fiore *et al*., 2008; Im and Kenny, 2012; Napoli and Pizzorusso, 2017;Rajman and Shartt, 2017; Cho *et al*., 2019].

Another important factor in neuroregeneration of pathological nervous system is that approximately 50% of the newborn neurons die because of the execution of programmed cell death [Oppenheim, 1991], even though during both development and adult neurogenesis the number of neurons that will die or mature varies considerably depending on the post-differentiation time, neuronal type, and CNS region [Pfisterer and Khodosevich, 2017]. Among all the mechanisms that prevent the new neuron from maturing is by directing it toward its apoptotic death [Fricker *et al*., 2018; Kuan *et al*., 2000]. The employment of multiple and redundant mechanisms to inhibit apoptosis enables mature neurons, with long-term survival capabilities, to become strikingly resistant to injury after the developmental stage [Kole *et al*., 2013; Hollvile *et al*., 2019]. Indeed, maturation of the CNS is associated with a decrease in the expression of several caspase genes [Kumar *et al*., 1992], including the effector caspase-3/7 [Kole *et al*., 2013].

Post-transcriptional regulation by microRNAs has been described among many modulators of apoptotic effectors (such as caspases) during the maturation of neuronal cells. Several microRNAs target the Casp3 gene (coding for caspase-3 or CASP-3 protein) and control its expression, such as miR-138-5p [Park *et al*., 2018;Maza *et al*., 2022]. This microRNA is one of the most enriched in the CNS [Obernosterer *et al*., 2006; Ludwig *et al*., 2016], mainly expressed in both brain and spinal cord neurons and varying depending on neuronal subtype [Maza *et al*., 2022]. It has heterogeneous expression and distribution throughout the neural tissue, which changes during CNS development[Obernosterer *et al*., 2006; Tatro *et al*., 2013; Daswani *et al*., 2022]. Moreover, it has also been shown that its expression levels change over the days of neuronal culture *in vitro* [Weiss *et al*., 2019]. On the other hand, miR-138-5p is known to regulate apoptotic cell death by regulating the expression of pro-apoptotic genes, among which Casp3 [Maza *et al*., 2022]. In the present work, we hypothesize that changes in miR-138-5p expression during neuronal maturation could control pro-apoptotic Caspase 3 expression and activity, thereby aiding the survival of neurons exposed to noxious stimuli.

## 2. Materials and Methods

### 2.1. Cell culture

We prepared primary cultures of hippocampal neurons from 18 days old (E18) Wistar rat embryos (RRID:RGD_13508588). After dissection from brain, we dissociated hippocampi by incubation with 1x trypsin (ThermoFisher) in Hanks’ Balanced Salt Solution (HBSS) medium without calcium and magnesium (Hyclone, GE Healthcare) supplemented with 20 mg/mL DNase (Roche) for 15 minutes at 37°C. We washed-out trypsin solution with HBSS with calcium and magnesium (Hyclone) before dissociating the tissue through repeated pipetting in Minimum Essential Medium (MEM; Gibco) supplemented with 10% horse serum (Fisher Scientific). We seeded the so-obtained cell suspension in 10 µg/mL poly-L-lysine (Sigma-Aldrich) pre-coated plates and let the cells adhere for 4 hours at 37°C and 5% CO_2_ in a cell culture incubator. Afterwards, we changed the medium to Neurobasal Medium (Gibco) enriched with 2% B-27 supplement (Gibco), 1% GlutaMAX (Gibco), and 100 u/mL penicillin/streptomycin (Gibco) and kept the culture in a humidified incubator in an atmosphere of 5% CO_2_ at 37°C for 1 or 14 days before subsequent experimental procedures.

### 2.2. Immunofluorescence assay

We seeded 35,000 hippocampal neurons per well in 24-well plates pre-coated with 10 µg/mL poly-L-lysine. One or 14 days later, we fixed cells with 4% PFA for 30 minutes and washed them with PBS 1x. Then, we permeabilized and blocked neurons in blocking buffer (3% BSA (Sigma-Aldrich) and 0,2% Triton X-100 diluted in PBS 1x buffer) for 1 hour at RT and incubated with antibody against β-III-tubulin (mouse anti-β-III-tubulin isoform, 1:500; Millipore cat#MAB1637, RRID: AB_2210524) and antibody against GFAP (chicken anti-GFAP, 1:1000; Abcam cat#ab4674, RRID: AB_304558) overnight (O/N) at 4°C. Antibodies were functionally validated by Millipore and Abcam companies, respectively. After incubation with primary antibodies, we washed cells with PBS 1x and incubated them for 1 hour at room temperature (RT) with a fluorescent Alexa 488-conjugated goat anti-mouse IgG secondary antibody (1:500; Molecular Probes cat#A-11029, RRID: AB_2534088) and a fluorescent Alexa 594-conjugated goat anti-chicken IgG secondary antibody (1:500; Molecular Probes cat#A-11042, RRID: AB_2534099). Finally, after three washes with PBS 1x, we mounted the coverslips on glass slides employing Fluorescence Mounting Medium (Thermo Scientific) with 1:30,000 of the fluorescent marker of nucleic acids 4’,6-diamino-2-fenilindol (DAPI; Sigma-Aldrich). We took photographs of the cells using an epifluorescence microscope (DMIL LED, Leica Microsystem GmbH) with a 20x microscope lens, coupled to a Leica DFC 3000G camera. We used the ImageJ software (National Institutes of Health, NIH) [Schindelin *et al*., 2012] to process and analyze the images.

We quantified the purity of hippocampal neuronal culture at 1 or 14 days of culture from the percentage of β-III-tubulin-stained neurons related to total cell number stained by DAPI (we analyzed a total of nine images per condition). On the other hand, we estimated the hippocampal neuronal density by calculating the total number of neurons per mm^2^ in the different cultures (we analyzed a total of five images of 0.27 mm^2^ per condition).

### 2.3. RT-qPCR analysis

We seeded 500,000 hippocampal neurons per well in 12-well plates with coverslips pre-coated with 50 µg/mL poly-L-lysine. One, 4, 7, 14 or 18 days later, we isolated and purified total RNA from neurons with miRNeasy Kit (Qiagen). For stimulated assays,, we treated 1 or 14 days primary cultures of hippocampal neurons O/N with 15 mM L-glutamic acid (LGA; Sigma-Aldrich). Total RNA concentration and purity (260/280 and 260/230 ratios) were estimated with a NanoDrop ND-1000 spectrophotometer (Thermo Scientific). Only samples with 260/280 ratios between 1.8 and 2.2 were employed.

To determine miR-138-5p expression, 10 ng of total RNA was reverse-transcribed and amplified using TaqMan microRNA gene expression assay (TaqMan® MicroRNA assay cat#002284, Applied Biosystems) following the manufacturer’s protocols. The U6 small nuclear RNA served as an internal control (TaqMan® MicroRNA assay cat#001973, Applied Biosystems). For mRNA detection of Casp3 transcripts, 1 µg of total RNA was subjected to random reverse transcription using Moloney Murine Leukemia Virus reverse transcriptase (M-MLV-RT; Invitrogen) and random primers (Roche). Then we evaluated the gene expression levels using TaqMan Gene Expression Assays for Casp3 (TaqMan® Gene Expression Assays cat#00563962; Applied Biosystems), employing 18S ribosomal RNA (TaqMan® Gene Expression Assays cat#4333760; Applied Biosystems) as a housekeeping gene. We measured the abundance of miR-138-5p and the mRNAs of interest in a thermocycler ABI Prism 7900 fast (Applied Biosystems) for 40 cycles of two steps: 15 seconds at 95°C plus 1 minute at 60°C using the 2^-ΔΔCt^ method [Livak and Schmittgen, 2001]. Briefly, the difference (ΔCt) between the cycle threshold of the microRNA or the target mRNA and their respective endogenous loading controls (U6 for miR-138-5p and 18S for Casp3) was estimated together with its associated variance following the standard propagation of error method from Headrick [Headrick, 2010]. Then, we compared the ΔCt value from different times of maturation with the ΔCt from 1 day to calculate the ΔΔCt and the correspondent fold increase (2^-ΔΔCt^), indicating also the 95% confidence interval (CI). In stimulated assays, we compared the ΔCt value from treated cells with the ΔCt from unstimulated cells (Ctr) to calculate the ΔΔCt and the correspondent fold change (2^-ΔΔCt^), indicating also the 95% confidence interval (CI).

### 2.4. Immunoblot assay

We analyzed pro-CASP-3 and cleaved-CASP-3 protein levels using standard immunoblot procedures. We seeded 500,000 hippocampal neurons per well in 12-well plates with coverslips pre-coated with 50 µg/mL poly-L-lysine. One or 14 days later, we stimulated a set of wells overnight with 15 mM LGA. After 24 hours, we incubated neuron lysates with radioimmunoprecipitation assay lysis buffer (RIPA, Sigma-Aldrich) containing a complete EDTA-free protease inhibitor cocktail (Roche) and centrifuged (14,000 g for 10 minutes at 4°C). Protein content was determined by the bicinchoninic acid method (BCA protein assay kit, ThermoFisher Scientific). We resolved a total of 50 µg of protein by SDS-PAGE, then electrophoretically transferred to a 0.2 µm polyvinylidene difluoride membrane (PVDF; Immobilon, Merck Millipore) and probed with antibody against CASP-3 (rabbit anti-CASP-3, 1:1000; Cell Signalling Technology cat#9662, RRID: AB_10694681) according to the manufacturer’s protocol. α-tubulin antibody was used as loading control (mouse anti-α-tubulin, 1:10000; Sigma-Aldrich cat#T6074, RRID: AB_477582). Antibodies were functionally validated by Cell Signalling Technology and Sigma-Aldrich companies, respectively. After incubation with the primary antibody, we washed membranes with TBS-Tween20 (Sigma Aldrich) and incubated for 1 hour at RT with a horseradish peroxidase (HRP)-conjugated goat anti-rabbit secondary antibody (1:1000; Thermo Fisher Scientific cat#31460, RRID: AB_228341) or an HRP-conjugated goat anti-mouse secondary antibody (1:1000; Thermo Fisher Scientific cat#31430, RRID: AB_228307). Finally, we developed HRP signal using the SuperSignal West Pico Chemiluminescent detection system (Pierce, ThermoFisher Scientific), and measured using ImageScanner III and LabScan v6.0 software (GE Healthcare Bio-Sciences AB) using default settings.

### 2.5. Measurement of Caspase-3/7 activity

To analyze effector caspase activity in stimulated hippocampal neurons we employed the CellEvent(tm) caspase-3/7 green detection assay (ThermoFisher Scientific) that allows analyzing activity in individual cells. Briefly, we seeded 35,000 hippocampal neurons per well in 24-well plates pre-coated with 10 µg/mL poly-L-lysine and, 1 or 14 days after, we stimulated cells O/N with 15 mM LGA. We assessed effector caspase activity 24 hours later by incubation in 2.5 μM of the assay reagent in warm PBS supplemented with 10% FBS medium for 30 minutes at 37°C protected from light. We took photographs of the cells using an epifluorescence microscope (DMIL LED) with a 20x microscope lens, coupled to a Leica DFC 3000G camera. We used ImageJ software to process and analyze the images. We estimated caspase-3/7 activity as the percentage of caspase-stained neurons related to the total neuron number.

### 2.6. Calcein/Propidium iodide assay

We seeded 35,000 hippocampal neurons per well in 24-well plates pre-coated with 10 µg/mL poly-L-lysine. One or 14 days later, we stimulated cells O/N with 15 mM LGA before incubating them with 2.5 µM calcein-AM (Sigma-Aldrich) and 0.4 µg/mL propidium iodide (PI; Sigma-Aldrich) in warm PBS supplemented with 10% FBS medium for 30 minutes at 37°C protected from light. We took photographs of the cells using an epifluorescence microscope (DMIL LED, Leica Microsystem GmbH) with a 20x microscope lens, coupled to a Leica DFC 3000G camera. We used the ImageJ software (National Institutes of Health, NIH) to process and analyze the images. Calcein-AM labels viable cells, whereas PI gains access only to cells with plasma membrane damage and accumulates in the nucleus. We estimated neuronal survival or death from, respectively, the percentage of calcein-AM or PI-stained cells related to the total neuron number.

### 2.7. Data analysis

Data are expressed as mean ± SEM as indicated in figure legends. Statistical significance of the treatment effects was tested using a two-tailed paired Student’s t-test or one-way or two-way ANOVA (followed by Tukeypost-hoc test for pairwise comparisons), depending on the characteristics of the data (the statistical analysis applied to each test is detailed in the figure legend).Statistical analyses and graphic representations were conducted using Prism Software 5 (GraphPad Software Inc.). Differences were considered statistically significant when the *p*-value was below or equal to 0.05.

## 3. Results

### 3.1. Opposite expression of miR-138-5p and Caspase 3 during hippocampal neuronal maturation

In order to establish the miR-138-5p expression along hippocampal neuronal maturation and to evaluate its possible effect on apoptotic proteins, we studied miR-138-5p levels in hippocampal neuron cultures during 18 days of maturation. RT-qPCR analyses show a significant upregulation of miR-138-5p at 7, 14 and 18 days of culture related to 1 day (*p*< 0.001, see Figure 1A and table associated for gene expression details; observe that ΔCt values are inversely related to gene expression). In the case of the expression of its target apoptotic gene of Caspase 3, we observed a significant decrease in Casp3 levels from 14 days of culture (*p*< 0.05, see Figure 1B and table associated). Besides, as shown in Figure 1C and D, immunoblot assays confirm the significant downregulation of pro-CASP-3 expression at 14 days of culture related to 1 day (14 d = 0.24% ± 0.04 v.s. 1 d = 0.85% ± 0.06; two-tailed paired t-test, T_3_ = 13.21, *p* = 0.0009).

**Figure 1.**
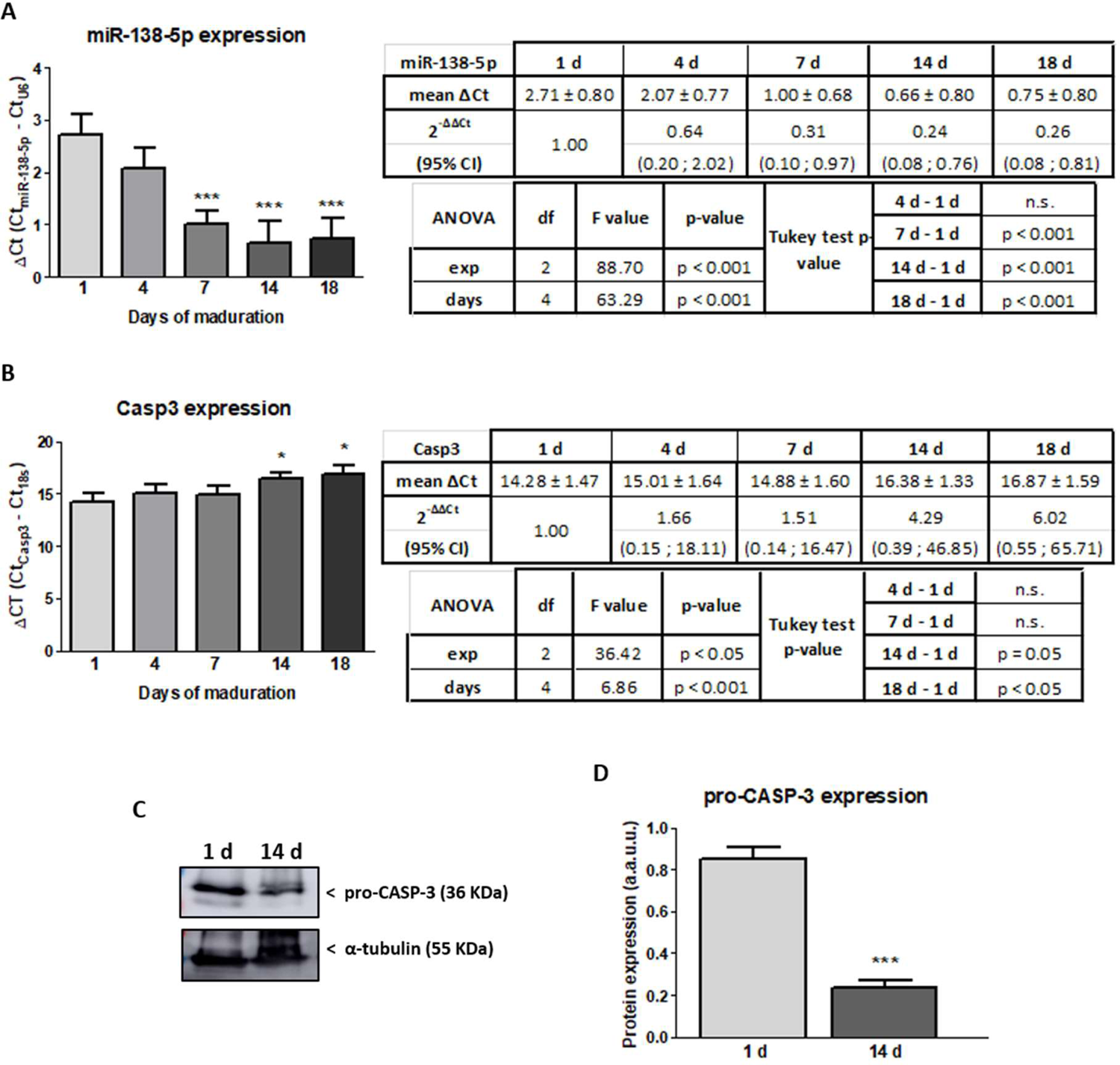
Endogenous expression levels of miR-138-5p and Caspase 3 in hippocampal neurons over maturation time. A) and B) RT-qPCR showing relative miR-138-5p (A) and Casp3 (B) expression in RNA isolated from hippocampal neuron cultures at 1, 4, 7, 14 and 18 days of maturation. Expression of each gene (Ct) was normalized to the Ct of its corresponding control gene (snoRNA U6 for miR-138-5p (ΔCt = Ct_miR138-5p_ - Ct_U6_) and 18S for Casp3 (ΔCt = Ct_Casp3_ - Ct_18S_)). Bars represent the mean ± SEM of four independent experiments. Tables show the microRNA and mRNA expression of miR-138-5p and Casp3 as the mean ΔCt and the fold change 2^-ΔΔCt^ (values and limits in folds of the 95% CI) of four independent experiments per time. Statistical analysis was carried out through two-way ANOVA and Tukey post-hoc tests. C) Representative immunoblot images of pro-CASP-3 and the load control α-tubulin protein expression in hippocampal neuron cultures at 1 or 14 days of maturation. D) Bars summarize the mean ± SEM of pro-CASP-3 protein expression of hippocampal neuron cultures at 1 or 14 days, normalized to the control protein expression α-tubulin of four independent experiments. Statistical analysis was carried out through a two-tailed paired t-test. * and *** denote significant differences: *p*< 0.05 and *p*< 0.001, respectively.

These results indicate that changes in expression of miR-138-5p and Caspase 3 are opposite after 14 days, so for the following experiments related to neuroprotection we select cells at 1 and 14 days of maturation.

### 3.2. Neuronal purity of hippocampal cell cultures

Before performing the different experiments with the cytotoxic treatment, we measured the purity of 1 and 14 days hippocampal neuronal cultures to avoid biases and correctly assign the neuronal effect in the following functional analysis. We observed that the percentage of neurons (β-III-tubulin-stained cells) at 1 day of culture related to total cells in culture was 82.02% ± 4.81. However, at 14 days this percentage decreased to 66.69% ± 5.20 (Figure 2). To avoid making a neuronal counting error due to this difference in purity over the days of culture, we only considered cells with a neuronal phenotype in the remaining assays performed in the single-cell analysis.

**Figure 2.**
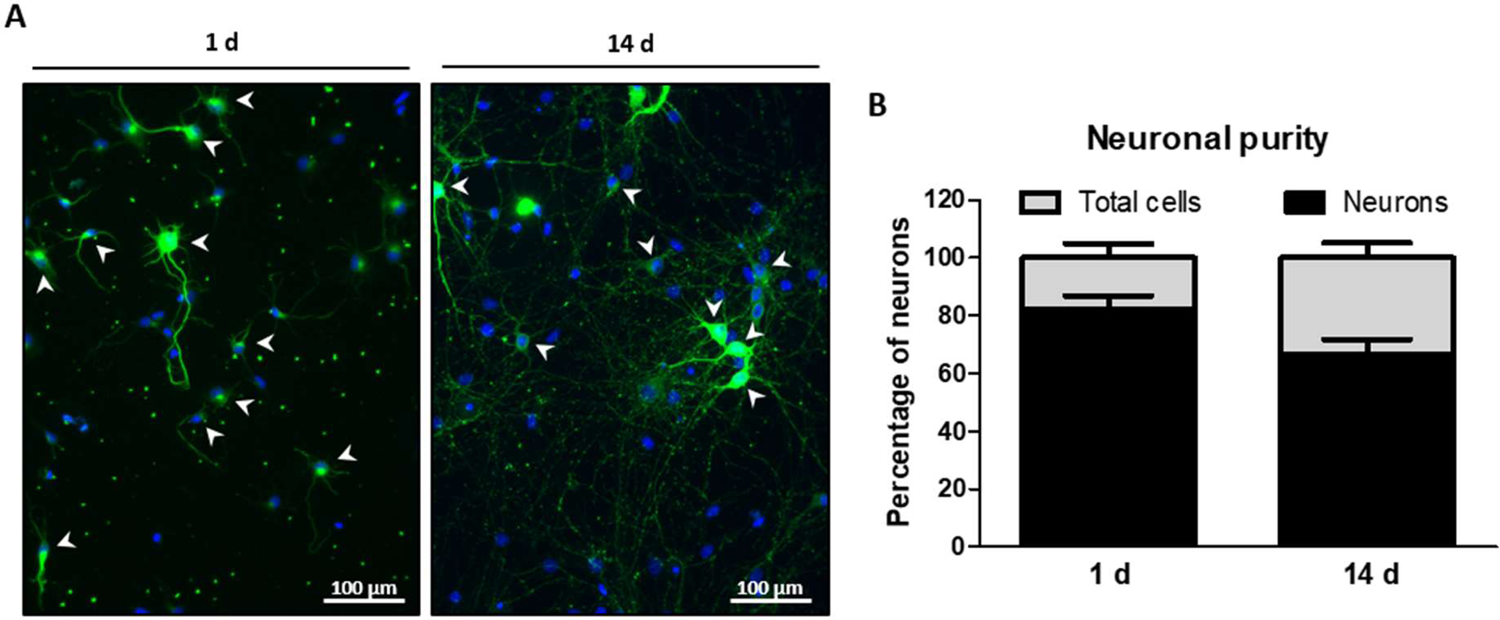
Purity of hippocampal neuronal cultures. Cell purity in hippocampal neurons at 1 or 14 days of maturation. A) Representative epifluorescence images of hippocampal neuron cultures at 1 and 14 days, labelled with the specific neuronal marker β-III-tubulin (green) and DAPI (nuclei staining, blue). Bar scale = 100 μm. B) Bar graph shows the mean of the percentage of neurons (β-III-tubulin-stained cells) related to total (DAPI positive) cells in culture ± SEM of five independent experiments.

### 3.3. Neuronal maturation attenuates caspase-dependent apoptosis

In order to evaluate injury response throughout days of maturation, we analyze the cellular levels of Caspase 3 in hippocampal neurons stimulated or not with LGA. We observed that treatment with LGA did not change the endogenous protein levels of pro-CASP-3 on neurons relative to untreated cells (Ctr) neither at 1 day of culture (LGA = 87.39% ± 8.923 v.s. Ctr (unstimulated after 1 day culture) = 100%; Figure 3A and B), nor at 14 days of maturation (LGA = 111.2% ± 8.319 v.s. Ctr (unstimulated after 14 days culture) = 100%; Figure 3A and B).

**Figure 3.**
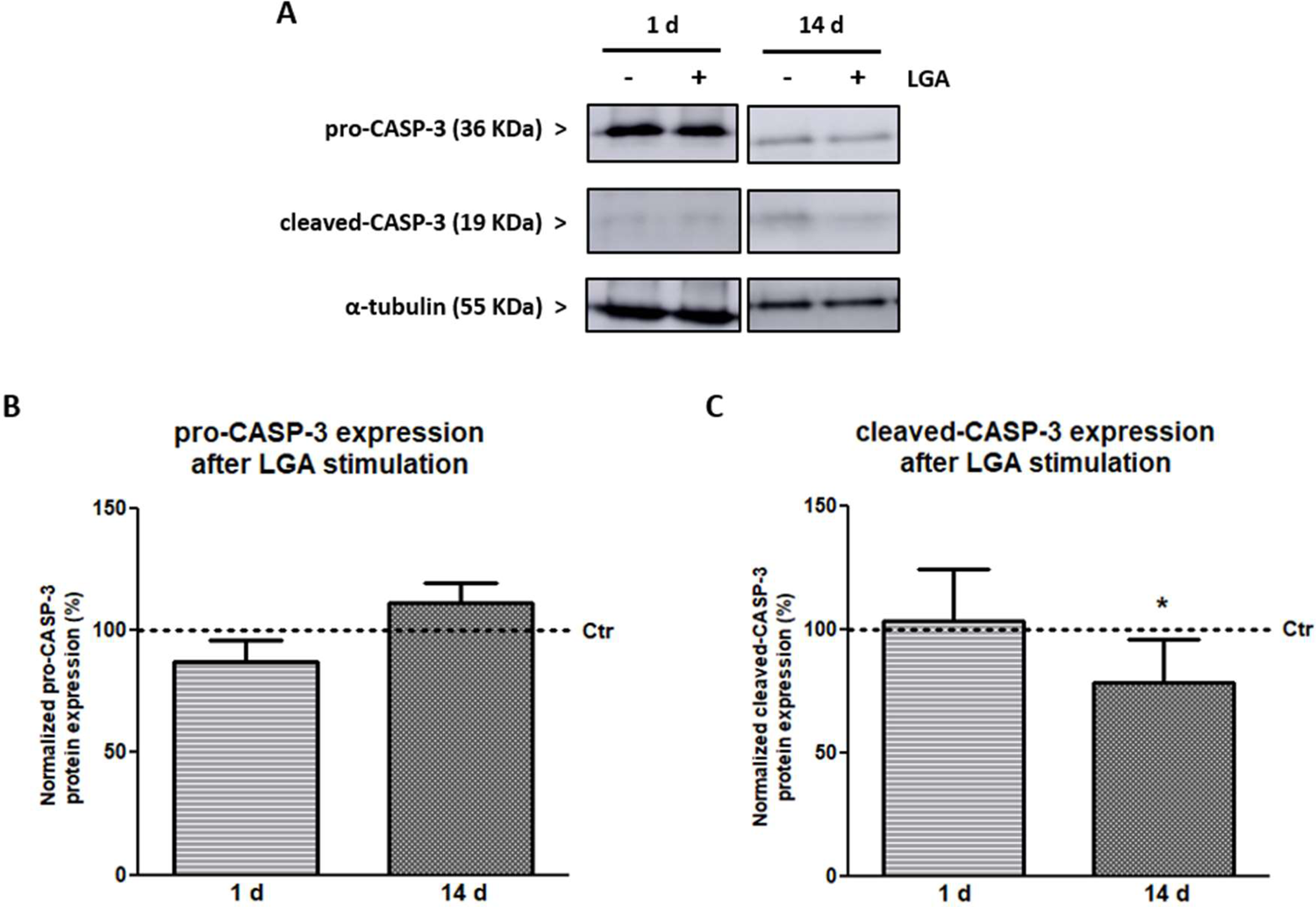
The expression of cleaved-CASP-3 is dismissed with neuronal maturation. Protein expression of pro-CASP-3 and cleaved-CASP-3 were analyzed in hippocampal neuron cultures stimulated with LGA at 1 or 14 days of maturation. A) Representative immunoblot images of pro-CASP-3, cleaved-CASP-3 and the load control α-tubulin protein expression in hippocampal neuron cultures treated with LGA or without treatment (Ctr) at 1 or 14 days. B) and C) Bars summarize the mean ± SEM of pro-CASP-3 and cleaved-CASP-3 expression after LGA treatment related to Ctr neurons of four independent experiments. Statistical analysis was carried out through a two-tailed paired t-test. * denotes a significant difference (*p*< 0.05).

A comparison of the protein expression levels of cleaved-CASP-3 in hippocampal neurons stimulated with LGA reveals that cleaved-CASP-3 expression did not change at 1 day of culture (LGA = 103.4% ± 21.34 v.s. Ctr (unstimulated after 1 day culture) = 100%; Figure 3A and C). However, at 14 days of maturation, the expression levels of cleaved-CASP-3 in neuronal cultures stimulated with LGA decreased related to unstimulated neurons (LGA = 78.49% ± 17.80 v.s. Ctr (unstimulated after 14 days culture) = 100%; Figure 3A and C). Thus, a significant change of cleaved-CASP-3 levels induced by LGA in neurons at 14 days of maturation relative to neurons at 1 day is observed (two-tailed paired t-test, T_2_ = 5.461, *p* = 0.032; Figure 3C).

### 3.4. Neuronal maturation reduces caspase activity

In order to establish the involvement of the pro-apoptotic caspase, we quantified the enzymatic activity of effector caspase-3/7 in hippocampal neurons after treatment. As shown in Figure 4A, at both 1 and 14 days, the activity of the effector caspase-3/7 of LGA treated cells increased related to unstimulated neurons (Ctr) (1 d: LGA = 204.4% ± 29.26 v.s. Ctr (unstimulated after 1 day culture) = 100%; 14 d: LGA = 120.3% ± 19.73 v.s. Ctr (unstimulated after 14 days culture) = 100%). We also tested whether maturation also reduced caspase-3/7 activity in cultures of hippocampal neurons after cytotoxic stimulation with LGA. The increase of enzymatic activity of neuronal effector caspase-3/7 induced by LGA is significantly lower in neurons at 14 days of culture than at 1 day (1 d = 1.044 ± 0.293, 14 d = 0.203 ± 0.197; two-tailed paired t-test, T_5_ = 4.265, *p* = 0.008; Figure 4B).

**Figure 4.**
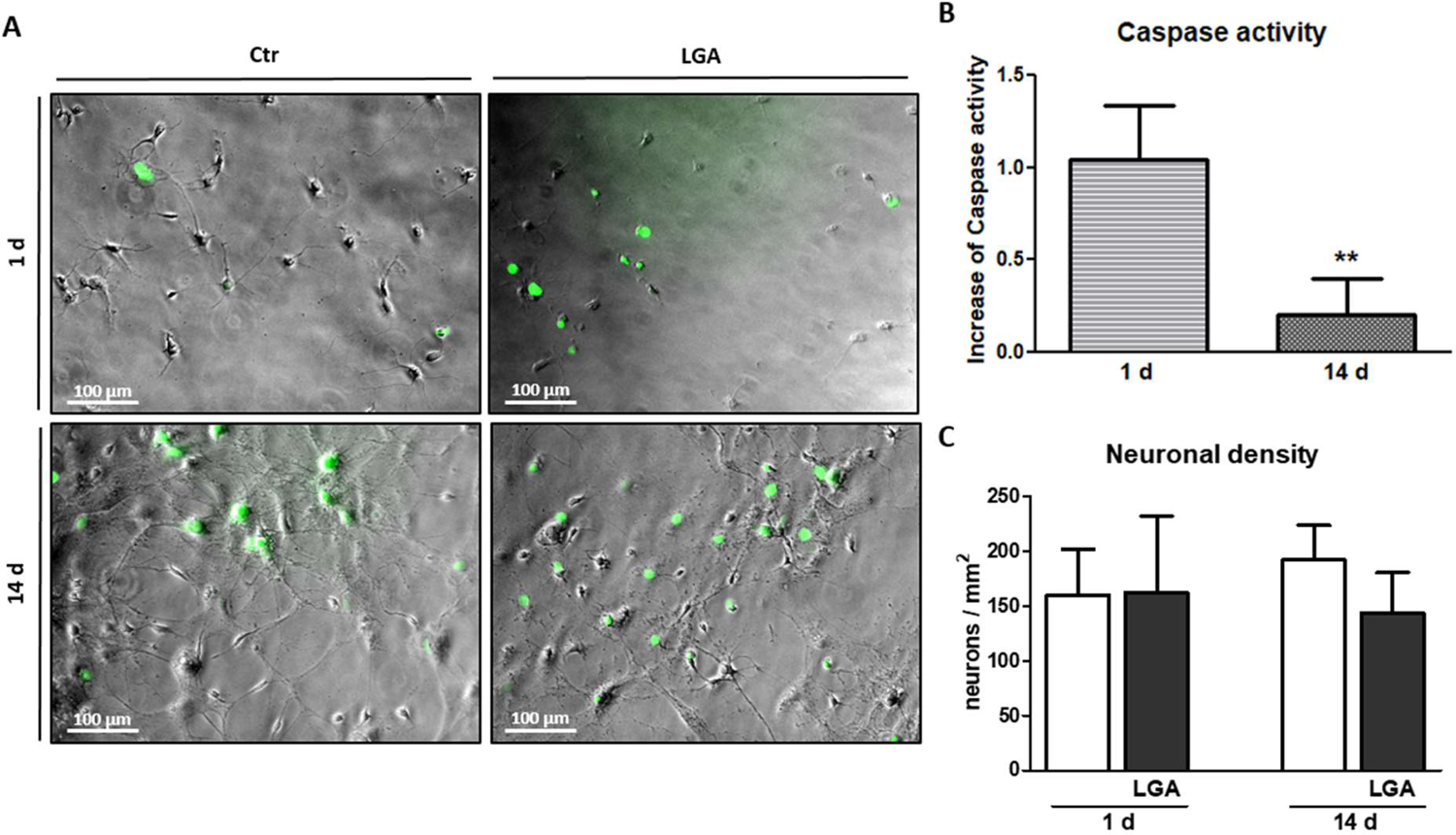
Neuronal maturation reduces caspase activity. Caspase activity in hippocampal neurons after LGA stimulation at 1 or 14 days of maturation. A) Representative phase contrast and epifluorescence images of hippocampal neuron cultures treated with LGA stimulus or without treatment (Ctr) at 1 or 14 days (green cells represent neurons stained with effector caspase reagent). Bar scale = 100 μm. B) Bars represent the mean ± SEM of the increase of activated effector caspase-stained neurons after LGA treatment related to Ctr neurons of six independent experiments. Statistical analysis was carried out through a two-tailed paired t-test. C) Bars summarize the mean ± SEM of the neuronal density (hippocampal neurons per mm^2^) of six independent experiments. Statistical analysis was carried out through one-way ANOVA and Tukey post-hoc tests. ** denotes significant differences (*p*< 0.01).

Despite observing different cell numbers at 1 and 14 days of culture, we did not observe differences in neuronal density (neurons per mm^2^) in the different culture conditions (maturation days or treatment) (1 d Ctr = 159.5 ± 42.36, LGA = 162 ± 70.08; 14 d Ctr = 192.1 ± 31.82, LGA= 143.5 ± 36.91) (Figure 4C).

### 3.5. Cell maturation promotes neuronal survival

To evaluate whether expression changes of cleaved-CASP-3 and caspase activity affect neuronal survival, we stimulated hippocampal neuron cultures with LGA at 1 and 14 days of culture and evaluated the percentage of live and dying cells in each sample. As shown in Figure 5, at both 1 and 14 days, when we treated cells with LGA neuronal survival (calcein-AM-stained cells) was reduced (1 d: LGA = 81.86% ± 3.377 v.s. Ctr (unstimulated after 1 day culture) = 100%; 14 d: LGA = 95.60% ± 5.217 v.s. Ctr (unstimulated after 14 days culture) = 100%) and cell death (PI-stained cells) increased (1 d: LGA = 209.2% ± 22.73 v.s. Ctr (unstimulated after 1 day culture) = 100%; 14 d: LGA = 113.3% ± 21.26 v.s. Ctr (unstimulated after 14 days culture) = 100%). On the other hand, and as shown in Figure 5A and B, the decrease of neuronal survival induced by stimulation with LGA is significantly lower in neurons at 14 days of culture than at 1 day (1 d = 0.181 ± 0.033, 14 d = 0.044 ± 0.052; two-tailed paired t-test, T_3_ = 4.422, *p* = 0.022). We confirmed the difference in cell death over the days of maturation using a PI assay. Complementary, hippocampal neuron cultures treated with LGA increased significantly less cell death at 14 days related to 1 day of culture (1 d = 1.092 ± 0.227, 14 d = 0.133 ± 0.217; two-tailed paired t-test, T_4_ = 4.227, *p* = 0.013; see Figure 5A and C).

**Figure 5.**
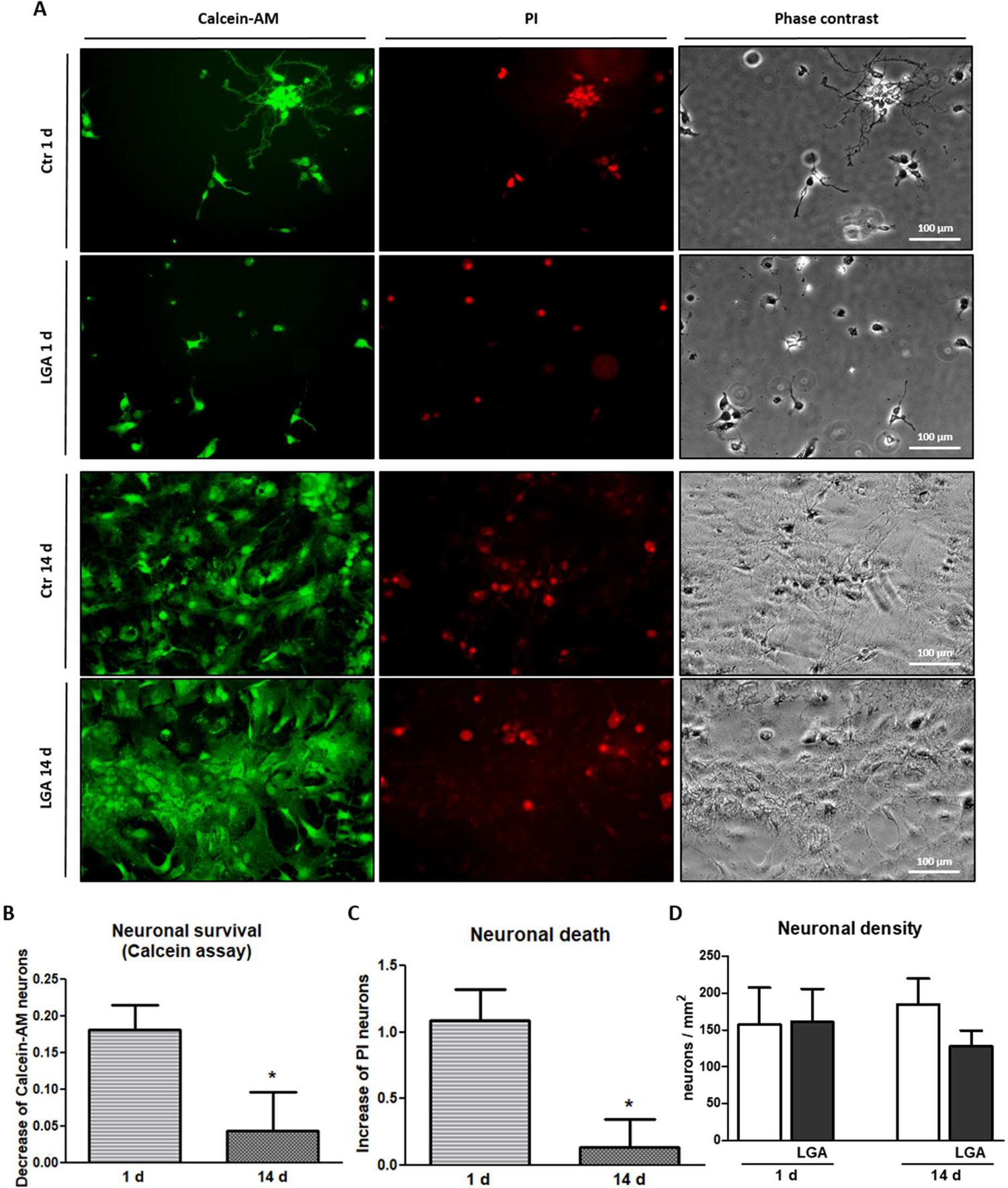
Neuronal maturation reduces cell death. Neuronal survival and death were analyzed in hippocampal neuron cultures stimulated with LGA at 1 or 14 days of maturation. A) Representative phase contrast and epifluorescence images of hippocampal neuron cultures treated with LGA stimulus or without treatment (Ctr) at 1 or 14 days (images of green cells represent cells stained with calcein-AM; images of red cells represent PI-stained cells; images in phase-contrast represent total cells). Bar scale = 100 μm. Bars summarize the mean ± SEM of the change in the percentage of neurons stained with calcein-AM (surviving neurons) (B) and PI (death neurons) (C) after LGA treatment related to Ctr neurons of five independent experiments. Statistical analysis was carried out through a two-tailed paired t-test. D) Bars represent the mean ± SEM of the neuronal density (hippocampal neurons per mm^2^) of five independent experiments. Statistical analysis was carried out through one-way ANOVA and Tukey post-hoc tests. * denotes significant differences (*p*< 0.05).

As in the caspase activity assay, we observed different number of total cells at 1 and 14 days of culture. However, we also did not observe any difference between different culture conditions in neuronal density in these cell survival or death assays (1 d Ctr = 157.3 ± 50.35, LGA = 160.9 ± 44.80; 14 d Ctr = 184.9 ± 34.96, LGA= 127.9 ± 21.13) (Figure 5D).

Altogether, these results indicate that cell maturation promotes neuronal survival in cytotoxic conditions.

### 3.6. Neuronal maturation does not change the increase of miR-138-5p expression in apoptotic stimulated hippocampal neurons

To evaluate whether the levels of the Casp3 targeted miR-138-5p change under cytotoxic conditions and to establish a possible relationship with apoptosis, we stimulated hippocampal neuron cultures with LGA at 1 and 14 days of culture and studied the endogenous expression of miR-138-5p. Initially, RT-qPCR analyses confirm the upregulation of miR-138-5p at 14 days related to 1 day of culture in endogenous conditions (Ctr) (14 d = 1.87 ± 0.29 v.s. 1 d = 2.57 ± 0.67; observe that ΔCt values are inversely related to gene expression), as well as after LGA treatment (14 d = 1.85 ± 0.11 v.s. 1 d = 2.81 ± 0.81). Besides, data show the significant downregulation of miR-138-5p after LGA stimulation at 1 day of culture (one-tailed paired t-test, T_3_ = 2.854, *p* = 0.05; Figure 6 and related table). However, at 14 days of maturation, the expression of miR-138-5p after LGA treatment did not change related to unstimulated neurons (Ctr; Figure 6 and related table). The comparison between both times shows a significant difference after treatment at 14 days of maturation related to 1 day (two-way ANOVA, F= 5.964, *p* = 0.040; see Figure 6). With these results, we proved that the increase of miR-138-5p expression throughout maturation does not change under apoptotic stimulation.

**Figure 6.**
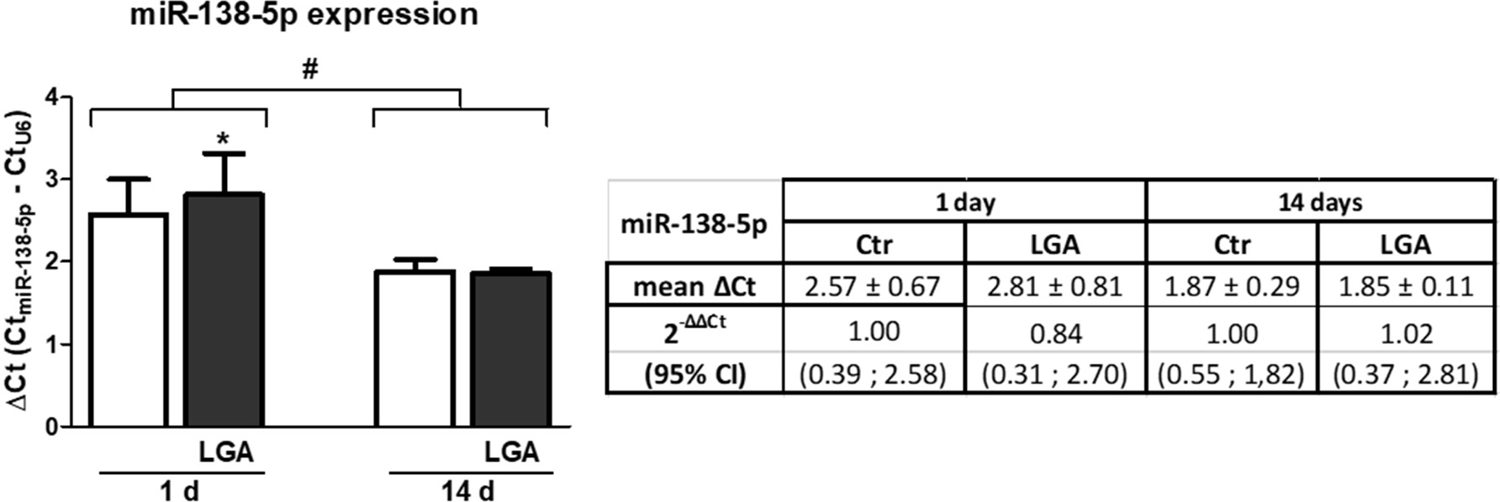
Differences in miR-138-5p expression after LGA stimulation at different days of maturation. RT-qPCR showing relative miR-138-5p expression in RNA isolated from hippocampal neuron cultures stimulated with LGA at 1 or 14 days of maturation. The expression of miR-138-5p (Ct) from each sample was normalized to the Ct of the control gene snoRNA U6 (ΔCt = Ct_miR138-5p_ -Ct_U6_). Bars summarize the mean ± SEM of three independent experiments. The table shows the expression of miR-138-5p as the mean ΔCt and the fold change 2^-ΔΔCt^ (values and limits in folds of the 95% CI) of three independent experiments per time. Statistical analysis was carried out through a one-tailed paired t-test and two-way ANOVA. * denotes significant differences (*p*<0.05) relative to unstimulated neurons. # denotes significant differences (*p* ≤ 0.05) between 1 and 14 days of neuronal cultures after LGA treatment.

## 4. Discussion

CNS development depends on a sculpting process that removes neural cells through programmed cell death. Apoptosis is widely acknowledged as the main regulator of cell number during development, which needs to be controlled in order to avoid neuronal overload and deleterious malformations in the nervous system [Buss *et al*., 2006; Elmore, 2007; Yamaguchi and Miura, 2015; Nguyen *et al*., 2021]. On the other hand, the modulation of apoptotic activation in newborn neurons upon pathological situations, such as injury or neurodegenerative diseases, is essential for neuronal maturation and tissue regeneration [Moujalled *et al*., 2021]. Thus, regulation of neuronal maturation is key to understanding not only how neurons survive throughout the life of an organism but also to shed light on how neuronal injury in neurodegenerative pathologies can lead to cell death. Among the numerous factors involved in determining cell fate during CNS maturation (e.g. cellular environment, growth factors, transcriptional regulation, etc.) [Belmonte-Mateos and Pujades, 2021], we focused on microRNA post-transcriptional regulation of pro-apoptotic factors. In this study, we observed an opposite pattern in the change of expression of endogenous Casp3 and miR-138-5p during hippocampal neuron maturation. Interestingly, miR-138-5p expression in mature neurons is not altered following apoptotic stimulation in contrast to the significant downregulation in immature neurons. Similarly, during neuronal maturation, not only the expression but also the processing of CASP-3 and caspase activity are reduced after apoptotic stimulation, resulting in a decrease of cell death.

Our results showing a decrease of Casp3 expression during maturation agreed with other studies showing that development of the nervous system is associated with decreased expression of Caspase 3 [de Bilbao *et al*., 1999] or other pro-apoptotic effectors, such as Caspase 7 or BAK [Zhang *et al*., 2007; Hollvile *et al*., 2019]. These changes are due, among other factors, to changes in their transcriptional regulation during neuronal development or maturation [Hollvile *et al*., 2019; Sousa and Flames, 2022]. The rate of transcription initiation of Casp3 substantially declines during brain maturation and is associated with transcriptional silencing by epigenetic regulation such as differential DNA methylation and histone acetylation [Yakovlev *et al*., 2010]. In addition to transcriptional regulation, post-transcriptional control of gene expression by microRNAs is crucial for the different phases of neuronal maturation [Rajman and Schratt, 2017]. Several microRNAs are key regulators of neuronal maturation, acting as inhibitors of neuronal apoptosis, such as miR-29, which modulates Bim, Bmf and Puma proteins expression [Kole *et al*., 2011; Swahari *et al*., 2021]. Similarly, miR-138-5p which is involved in migration, axonal growth or in determining the size of dendritic spines [Kisliouk and Meiri, 2013; Liu *et al*., 2013; Bicker *et al*., 2014], has been shown to regulate both the expression and activity of pro-apoptotic factors (i.e. Casp3, Casp7 and Bak1) following traumatic spinal cord injury [Maza *et al*., 2022]. This microRNA is upregulated during CNS maturation as observed previously in cortical development [Liu *et al*., 2013]. From our results, we observed an increased expression level of miR-138-5p in cultured hippocampal neurons during maturation, described before in cortical neuron cultures by Weiss and cols. [Weiss *et al*., 2019]. These expression changes during cellular maturation is not restricted only to neurons but also occurs in other cell types, being upregulated in oligodendrocytes, promoting the early stages of maturation and facilitating the appropriated axonal myelination [Dugas *et al*., 2010]. Thus, although maturation is promoted in both cases, the function of the miR-138-5p overexpression could be cell-dependent.

Interestingly, upon exposure to a challenging cellular environment, the expression of these post-transcriptional regulators is also differentially altered depending on the neural maturation stage. The expression of miR-29a or miR-124 is altered during the maturation of neural cells in noxious conditions [Li e*t al*., 2013; Nampoothiri and Rajanikant 2017]. The Casp3 regulator miR-29b is downregulated in immature cerebellar neurons after ethanol exposure and recovers its high expression levels during the following days of culture, thus protecting cells from apoptotic activation [Qi *et al*., 2014]. Similarly, we have shown that the expression of miR-138-5p in mature neurons is not altered under cytotoxic conditions (LGA stimulus), though a significant downregulation is observed in young neurons. So, our results indicate that the distinct neuronal death sensibility according to its stage of maturation might depend on the altered expression of miR-138-5p and many other microRNAs.

Our results on neuronal maturation showed the attenuation of caspase-dependent apoptosis and the promotion of neuronal survival. We observed, studying hippocampal neurons individually, that the number of neurons with effector caspase-3/7 activity decreased significantly under cytotoxic conditions (LGA stimulus) in mature neurons. Similarly, it has been shown age-dependent differences in both injury-induced Caspase 3 activation and susceptibility to apoptosis in the mammalian brain [Bittigau *et al*., 1999; Pohl *et al*., 1999; Hu *et al*., 2000], as well as a reduction of etoposide-induced apoptosis and decreased levels of Caspase 3 activity in primary rat cortical neurons [Yakovlev *et al*., 2001]. Caspase-3/7 effector activity has different levels of regulation at the transcription, post-transcription and translation stages. In order to post-transcriptionally control the expression of the pro-apoptotic factors, such as Casp3, during the different phases of neuronal maturation, it has been considered microRNA regulation, differential subcellular compartmentalization of specific effector caspases, and changes in the expression of direct regulators of Caspase 3 activity. The spatial confinement of active Caspase 3 -where the control of its activity may be different in the soma than in the dendrites [Ertürk *et al*., 2014]-has been described since it is also involved in axon guidance, synapse formation, axon pruning, and synaptic functions aside from their roles in eliminating unnecessary neural cells [Nguyen *et al*., 2021]. On the other hand, the attenuation of Caspase 3 activation during maturation may result from the repression of factors involved in the Caspase 3 activation pathway, as the observed reduction of cytochrome c ability to induce activation of Casp3 in the murine brain, being undetectable after 2 weeks of age [Yakovlev *et al*., 2001]. However, other regulators, such as the X-linked inhibitor of apoptosis protein (XIAP), have been shown to decrease its levels in the cerebral cortex both in ageing and in cultured cortical neurons [Fadó *et al*., 2013].

Therefore, our results point out that the observed opposite expression of endogenous Casp3 and miR-138-5p might influence controlling apoptosis, and favouring neuronal survival upon cell death stimulation at distinct neuronal maturation stages. However, further study would be needed to determine a more detailed role of miR-138-5p in apoptosis during neuronal maturation using loss-of-function assays with the downregulation of the microRNA using antagomiRs. Moreover, this regulation of apoptosis could be due not only to the action of miR-138-5p but also to the synergistic action of several microRNAs acting cooperatively on Casp3 or other targets in the apoptotic pathway. In conclusion, our results provide a broader role of miR-138-5p to its already widely studied involvement in other pathologies, such as cancer. We describe its important implication concerning neuronal maturation, regulating Caspase 3 and favouring survival, which is essential in the creation of new neurons in injury or neurodegenerative diseases.

## Supporting information

Original WB images

## 5. Declaration Section

### Statement of Ethics

All experimental procedures were in accordance with the European Communities Council Directive 2010/63/EU, Spanish Royal Decree 53/2013 (experimental animal use regulation) and Order ECC/566/2015 (regulation of personnel formation in animal experimentation) and were approved by the Hospital Nacional de Parapléjicos Animal Care and Use Committee (project ref #153BCEEA/2016).

### Disclosure Statement

The authors have no conflicts of interest to declare.

### Funding Sources

The research was funded by grants from the Fundación Tatiana Pérez de Guzmán el Bueno and the Council of Education, Culture and Sports of the Regional Government of Castilla La-Mancha (Spain) and Co-financed by the European Union (FEDER) “A way to make Europe” (project references SBPLY/17/000376 and SBPLY/21/180501/000097). M. Asunción Barreda-Manso was funded by the Council of Health of the Regional Government of Castilla La-Mancha (Spain), through the “Convocatoria de Ayudas Regionales a la Investigación en Biomedicina y Ciencias de la Salud”, II-2020_05. Altea Soto was funded by the Council of Education, Culture and Sports of the Regional Government of Castilla La-Mancha (Spain).

### Author Contributions

MABM, RMM and MND were responsible for the conception and design of the experiments. MABM was responsible for the acquisition and analysis of the data. MABM and RMM were responsible for the interpretation of the data and to draft and revise the manuscripts.AScontributed to the acquisition of the data. AS, TMG, DR and MND contributed to the revision of the manuscripts.MABM and RMM were responsible for drafting and revising the final version of the manuscript. RMM and MND were responsible for funding acquisition.All authors have revised and approved the final version of this article and express their agreement for all aspects of the present work in ensuring that questions related to the accuracy or integrity of any part of the work are appropriately investigated and resolved. All authors read and approved the final manuscript.

## Acknowledgements

We thank the Fundación del Hospital Nacional de Parapléjicos para la Investigación y la Integración (FUHNPAIIN) and theanimal facility from the Research Unit of the Hospital Nacional de Parapléjicos (Toledo, Spain) for their technical and logistic support.

## Notes

### Competing Interest Statement

The authors have declared no competing interest.

### Summary of Updates

Author names and affiliations updated; better image quality

