## Supplementary material for "MiR-138-5p upregulation during neuronal maturation parallels with an increase in neuronal survival": Original WB images

**Figure 1: Raw immunoblot images of pro-CASP-3 in hippocampal neurons at 1 or 14 days of maturation.** Red rectangles surround the samples used for statistical analysis. Green rectangles surround the samples used to illustrate the figure. Exp 1 to 4 indicate the experiment number. 1 d indicates neurons of 1 day of culture whereas 14 d codes for neurons of 14 days of maturation.

**Figure 1.C**

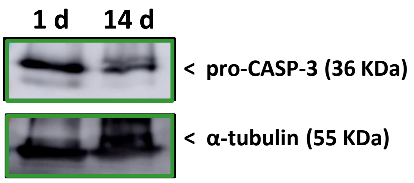


**Raw images**

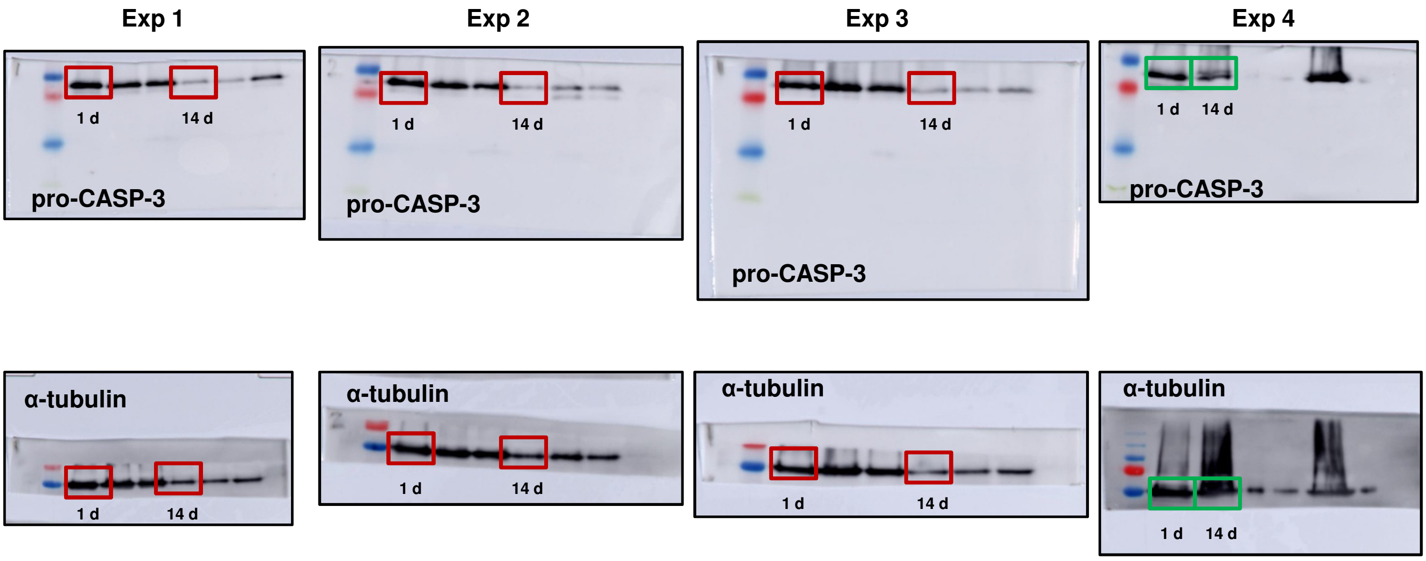

**Figure 3: Raw immunoblot images of pro-CASP-3 and cleaved-CASP-3 in hippocampal neurons stimulated with LGA at 1 or 14 days of maturation.** Red rectangles surround the samples used for statistical analysis. Green rectangles surround the samples used to illustrate the figure. Exp 1 to 4 indicate the experiment number. 1 d indicates neurons of 1 day of culture whereas 14 d codes for neurons of 14 days of maturation. LGA indicates neurons treated with LGA whereas C codes neurons without any treatment.

Figure 3.A

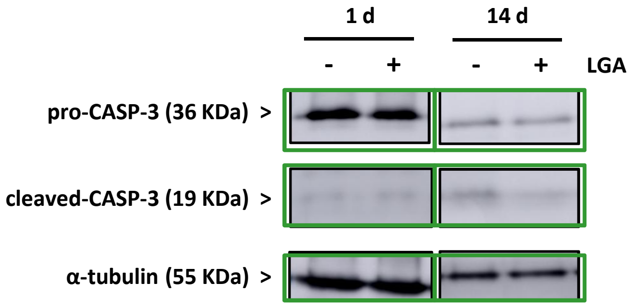


Raw images

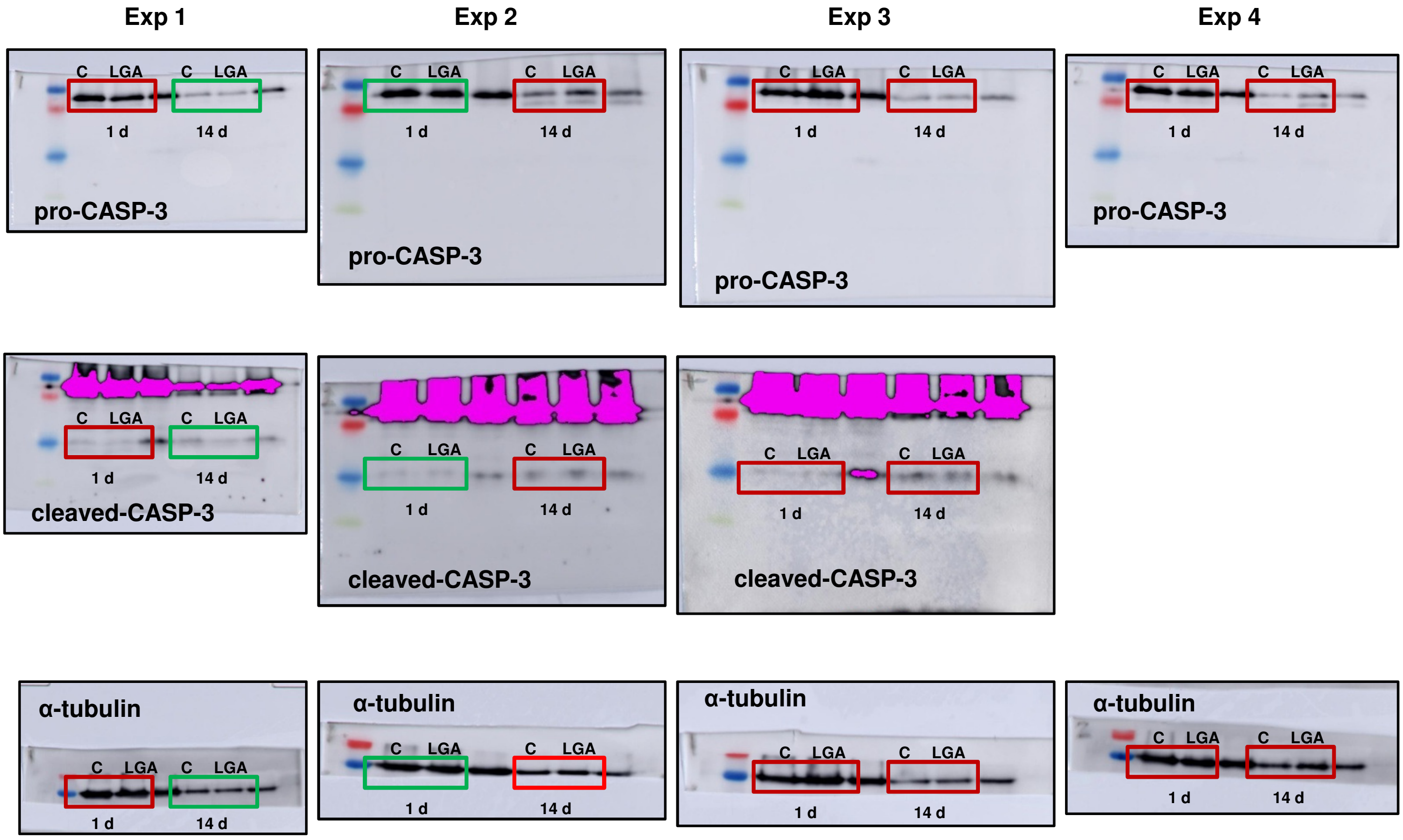
